# Developing a Novel Positronium Biomarker for Cardiac Myxoma Imaging

**DOI:** 10.1101/2021.08.05.455285

**Authors:** Paweł Moskal, Ewelina Kubicz, Grzegorz Grudzień, Eryk Czerwiński, Kamil Dulski, Bartosz Leszczyński, Szymon Niedźwiecki, Ewa Ł. Stępień

## Abstract

Here, positronium imaging is presented to determine cardiac myxoma (CM) extracted from patients undergoing urgent cardiac surgery due to unexpected atrial masses. Positronium is an atom build from an electron and a positron, produced copiously in intra-molecular voids during the PET imaging. CM, the most common cardiac tumor in adults, accounts for 50-75% of benign cardiac tumors. We aimed to assess if positronium serves as a biomarker for diagnosing CM. Perioperative examinations and histopathology staining in six patients confirmed the primary diagnosis of CM. We observed significant differences in the mean positronium lifetime between tumor and normal tissues, with an average value of 1.92(02) ns and 2.72(05) ns for CM and the adipose tissue, respectively. Our findings, combined with positronium lifetime imaging, reveals the novel emerging positronium biomarker for cardiovascular imaging.

**One-Sentence Summary:** Positronium may serve as an imaging biomarker for cancer diagnostics.

---

The advent of total-body positron emission tomography (TB-PET) has opened a new paradigm for personalized care in precision medicine, enabling simultaneous kinetic and parametric molecular imaging of all tissues in the human body (*1-8*). The long axial field of view (covering the complete human body) and high sensitivity of TB-PET (40 times higher than standard PET scanners) enable the dynamic total-body scan at one bed position. This in turn facilitates quantitative diagnostic analyses based on parametric images, for example, the influx rate of radiopharmaceutical uptake, in addition to the conventional semi-quantitative standardized uptake value (SUV) based diagnostics (*9,10*). Moreover, the high sensitivity of TB-PET has developed new prospects for introducing in medicine novel diagnostic parameters based on positronium atoms, which are copiously produced in the human body during regular PET imaging (*1,2,11-13*).

Positronium is an exotic atom created from an electron (present in biomolecules) and a positron emitted by a radionuclide while being a part of a radiopharmaceutical. Figure 1 outlines the basic processes that lead to the formation and decay of positronium in the hemoglobin molecule. In PET, as much as 40% of positron-electron annihilations occurs through the production of positronium atoms in the inter- and intra-molecular spaces (voids) inside a patient’s body (*1,2,11-13*). The properties of these positronium atoms, such as lifetime and decay rate, depend on the size of molecular voids as well as the concentrations of biomolecules in tissues and biofluids. Moreover, they could provide information about the disease status or tumor microenvironment for discrimination a hypoxic region (*2,14*). In previous studies, we hypothesized that positronium imaging capabilities combined with TB-PET systems may lead to significant improvements in diagnostic and prognostic assessments (*1,2,11-13*). Total body approach will bring an advantage in the early diagnosis and treatment of various diseases (*8*), including oncological (*15*), cardiovascular (*16*), systemic and neurological (*17-18*), thus enabling the simultaneous detection of pathologies on a molecular level, before they lead to functional or structural abnormalities (*1*).

**Fig. 1.**
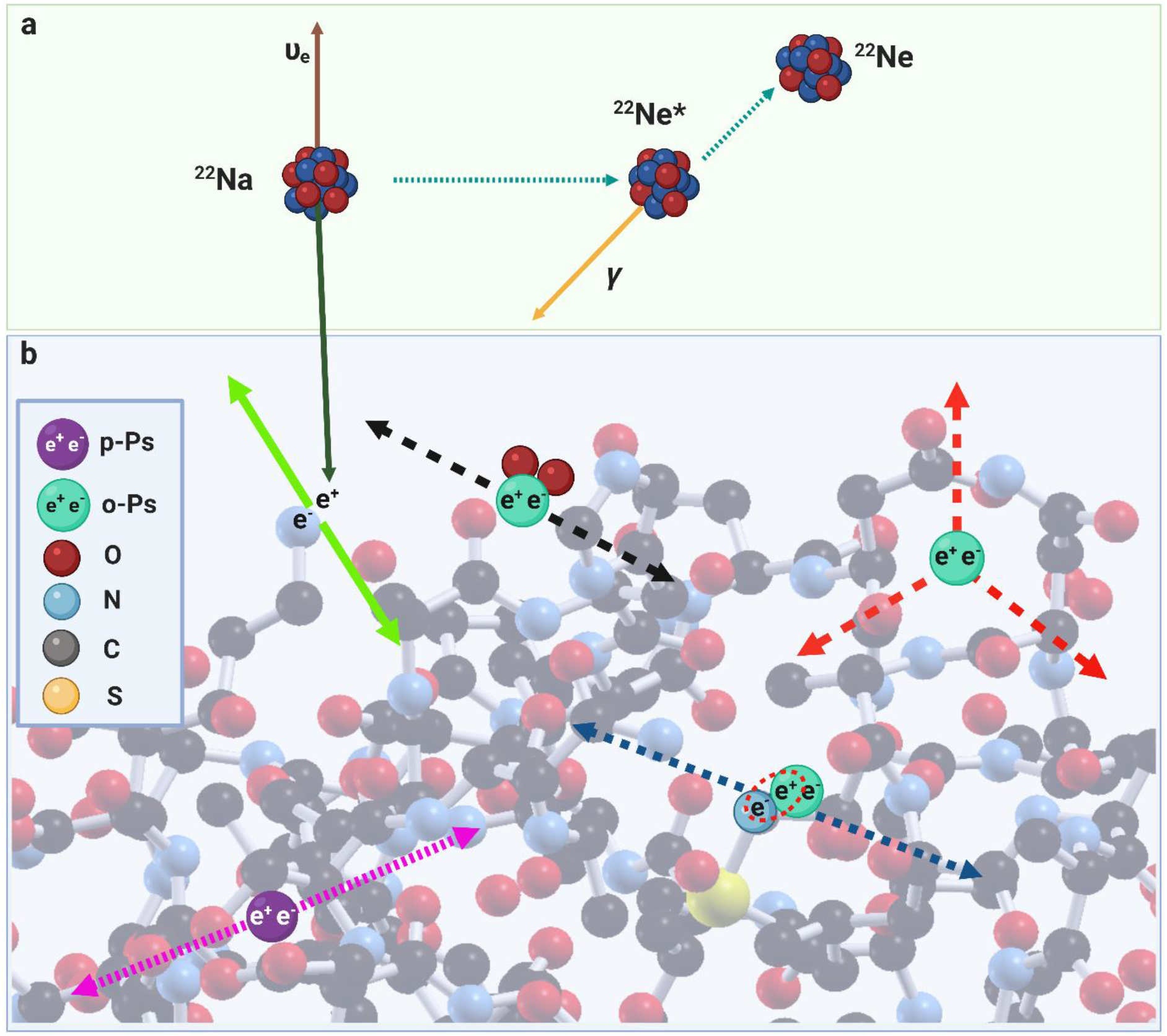
A pictorial illustration of the basic processes leading to the formation and decay of positronium in the intramolecular voids of a hemoglobin molecule. We have used a sodium ^**22**^Na isotope as an emitter of positrons (e+), considering the role of sodium fluoride in cardiovascular imaging (*32,35*). (A): ^**22**^Na radionuclide decays emitting a neutrino (brown arrow) and a positron (dark green arrow) (e+), and turns into an excited ^**22**^Ne* nucleus. It de-excites almost instantly (on an average in 3 ps) by the emission of the prompt photon (yellow arrow). (B): The positron thermalizes at a distance of about 1 mm (*36*), and annihilates into photons with one of the electrons (e^-^) in the surrounding molecules. Positron-electron annihilation in the tissue undergoes direct annihilation into two photons (solid light green arrows) in roughly 60% of the cases. However, it proceeds *via* the formation of positronium atom in 40% of the cases. The latter may be trapped in the tissue in the intra-molecular voids (*37*). Positronium atom can be created in two forms: (i) short-lived (125 ps) para-Positronium (p-Ps indicated in purple), which decays into two photons (dotted pink arrows) or (ii) long-lived (142 ns) ortho-Positronium (o-Ps indicated in mint), which decays into three photons (dashed red arrows). In the tissue, o-Ps predominantly annihilates either through an interaction with an electron (e^-^) from the surrounding molecule *via* pick-off process (dashed blue arrows) or through the conversion to p-Ps *via* an interaction with oxygen molecules, which subsequently decays into two photons (dashed black arrows) (*14,38*). These processes decrease the o-Ps lifetime, which becomes strongly dependent on the size of intramolecular voids and the concentration of bio-active molecules.

In this article, we demonstrate the strength of novel positronium diagnostic parameters for the in vivo assessment of benign cardiac tumors. We measure and analyze the mean positronium lifetime in a cardiac neoplasm (myxoma) and compare it with data obtained from normal mediastinal adipose tissues.

Cardiac myxoma (CM) was first described by Polish pathologist W. L. Brodowski in 1867 as a walnut size jelly-like tumor dissected from the left atrium of a 40-year-old woman who died of parenchymal nephritis (*19*). The first description of five left ancient ventricular polypus specimens was summarized by T. W. King in 1845 (20). CM is the most common cardiac tumor in adults and accounts for approximately 50-75% and >50% of benign and primary cardiac neoplasms, respectively (*21-26*). Left CM is twice most common in women, compared to men (*27*). Considering its left-atrial localization, CM is an origin of emboli to the vascular tree, particularly to the central nervous system (*27*). It results in stroke or transient ischemic attack, particularly in young adults (*28,29*). Therefore, CM is a life-treating disease that increases the risk of severe systemic and cardiac symptoms, perioperative morbidity, and mortality (*27*). The etiology of CM is poorly understood. Nevertheless, researchers have confirmed the presence of cardiomyocytes and mesenchymal progenitor factors (Nkx2.5/Csx, c-kit) in CM cells, thus revealing the stem cell origin of myxoma-initiating cells (*30,31*).

The diagnosis of CM is often elusive, especially in young stroke survivors (*29*). This can be attributed to their non-specific neurological symptoms. Transthoracic echocardiography (TTE) is the initial technique for the differential diagnostics of CM (*21,26,27*). It provides additional information about the hemodynamic parameters and enables the characterization of intracardiac masses, in terms of their size and morphology. However, TTE fails to distinguish between thrombi, vegetation, or neoplasms (*21*). In practice, cardiac computed tomography (CT) and magnetic resonance imaging (MRI) are less available for the majority of patients. Furthermore, the pre-operative imaging diagnosis appears to be discordant with histopathologic findings in one-fifth and more than half cardiac tumors and malignant lesions, respectively. This in turn limits CM recognition.

PET is used as standard in cardiovascular imaging, for example, for semi-quantitative myocardial perfusion evaluation (*32,33*). For research an assessment of atherosclerosis (16,34) or the diagnosis and risk stratification in patients with known or suspected coronary artery disease (*32,35*) Nevertheless, standard PET does not reveal the tissue histology and fails to determine if the organic cause of the detected disorder is cancer origin, inflammation, or intracardiac thrombus.

Herein, we first present the results indicating that the in vivo imaging of positronium properties, available because of TB-PET scanners (*1,2,11-13*), would facilitate the quantitative in vivo assessment of the intracardiac masses. Our findings would shift imaging diagnostics towards a new direction of personalized medicine.

## Results

### CM tumors obtained from symptomatic patients were localized in a left atrium and they were characterized by different morphology

We examined six patients, comprising four women. Their age ranged between 52 and 84 years. Moreover, they had multiple comorbidities, including metabolic and cardiovascular (table S1) conditions. For each patient, we conducted a screening interview and laboratory test (blood counts, metabolic panel, and urine test), and diagnosed them based on the transthoracic TTE examination. The patients were symptomatic and had TTE-confirmed cardiac masses, localized in a left atrium. This necessitated the cardiac surgery (Fig. 2A). We obtained perioperative specimens of CM and mediastinal adipose tissues from the operating theatre (John Paul II Hospital, Kraków). Half of each sample was fixed and designated for a histopathological analysis (Fig. 2A). In contrast, the other half was used for studying their positronium properties using PALS both in tissues and cell culture (Fig. 2B, D) and micro-CT examination (Fig. 2C).

**Fig. 2.**
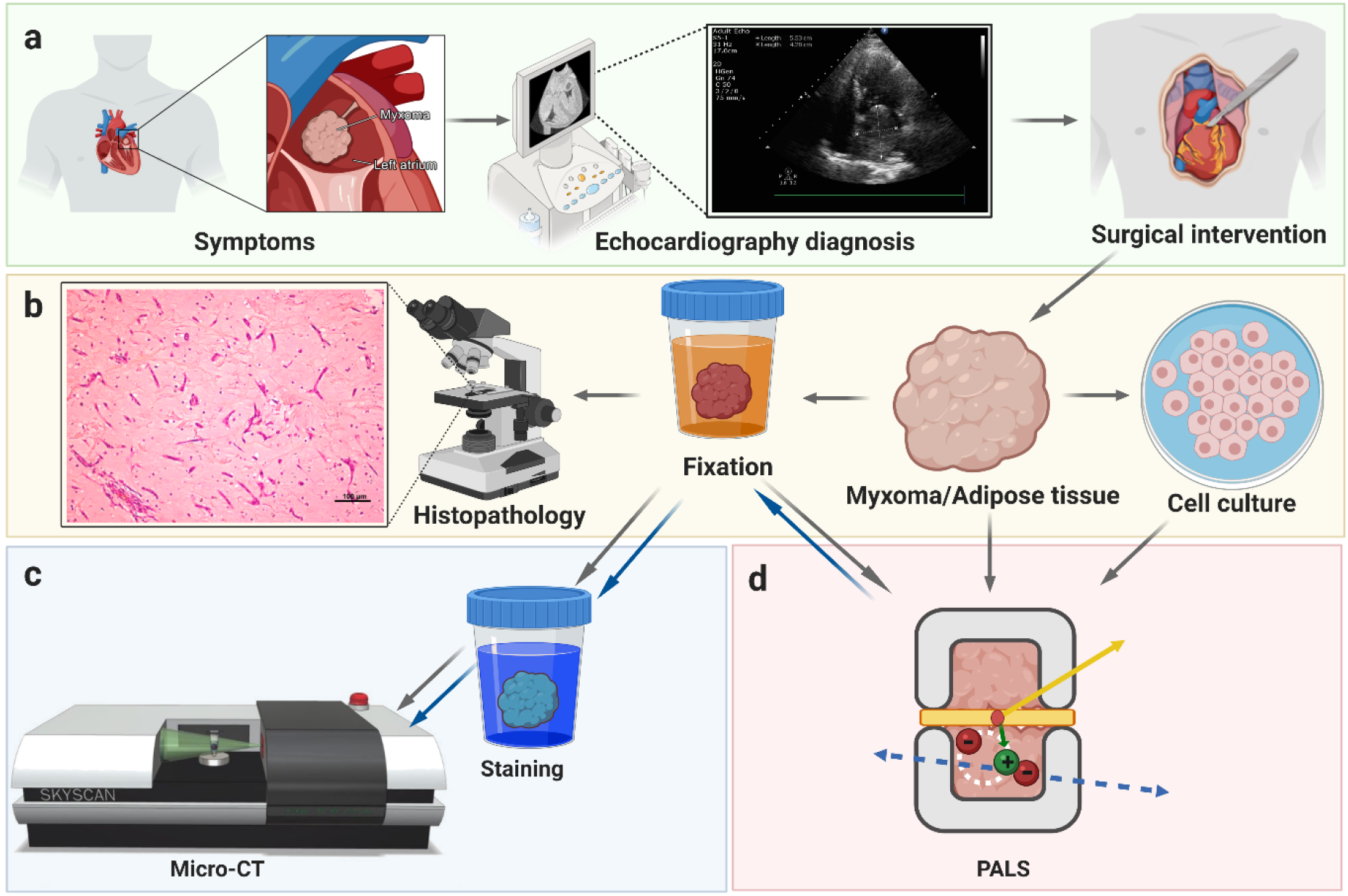
Cardiac myxoma experiment workflow. (A): The clinical examination of a symptomatic patient by transthoracic echocardiography and surgical excision of cardiac myxoma (CM) (B): Sample preparation and examination of myxoma, adipose tissues and CM cells: one piece of CM was fixed for histopathology examination (H&E), while other piece of CM and adipose tissue samples have been used for studying their positronium properties and later fixed for the X-ray imaging (micro-CT). CM cell culture has been derived from the part of tissue designated for PALS experiment before fixation. (C): Micro-CT imaging: staining in Lugol solution for 5 days and micro-CT scanning; gray and blue arrows represent a workflow for two different scanning runs: before and after positronium measurement (PALS). (D): A chamber for ortho-Positronium lifetime measurement comprising CM or adipose sample with the ^**22**^Na radionuclide (red dot) emits (green arrow) a positron (+) and a prompt photon (yellow arrow). Positron and electron from a sample create a positronium atom (bound state of electron and positron), indicated pictorially by a white dotted circle. An example annihilation of positronium into two photons (blue arrows). Fig. 3 contains a detailed description of positronium lifetime measurement.

Perioperative examinations and routine histopathology staining confirmed the primary diagnosis of CM. All cardiac tumors were pedunculated. They were diverse in size and structure. While four tumors were solid, two were of papillary type (table S1). In histopathology, we observed a typical hematoxylin and eosin (H&E) picture of a myxoma tumor with purple stellated or globularly shaped CM cells. Moreover, the orange-reddish structures represented the blood vessels with erythrocytes and the surrounding myxoid matrix was stained in pink (Fig. 3A).

**Fig. 3.**
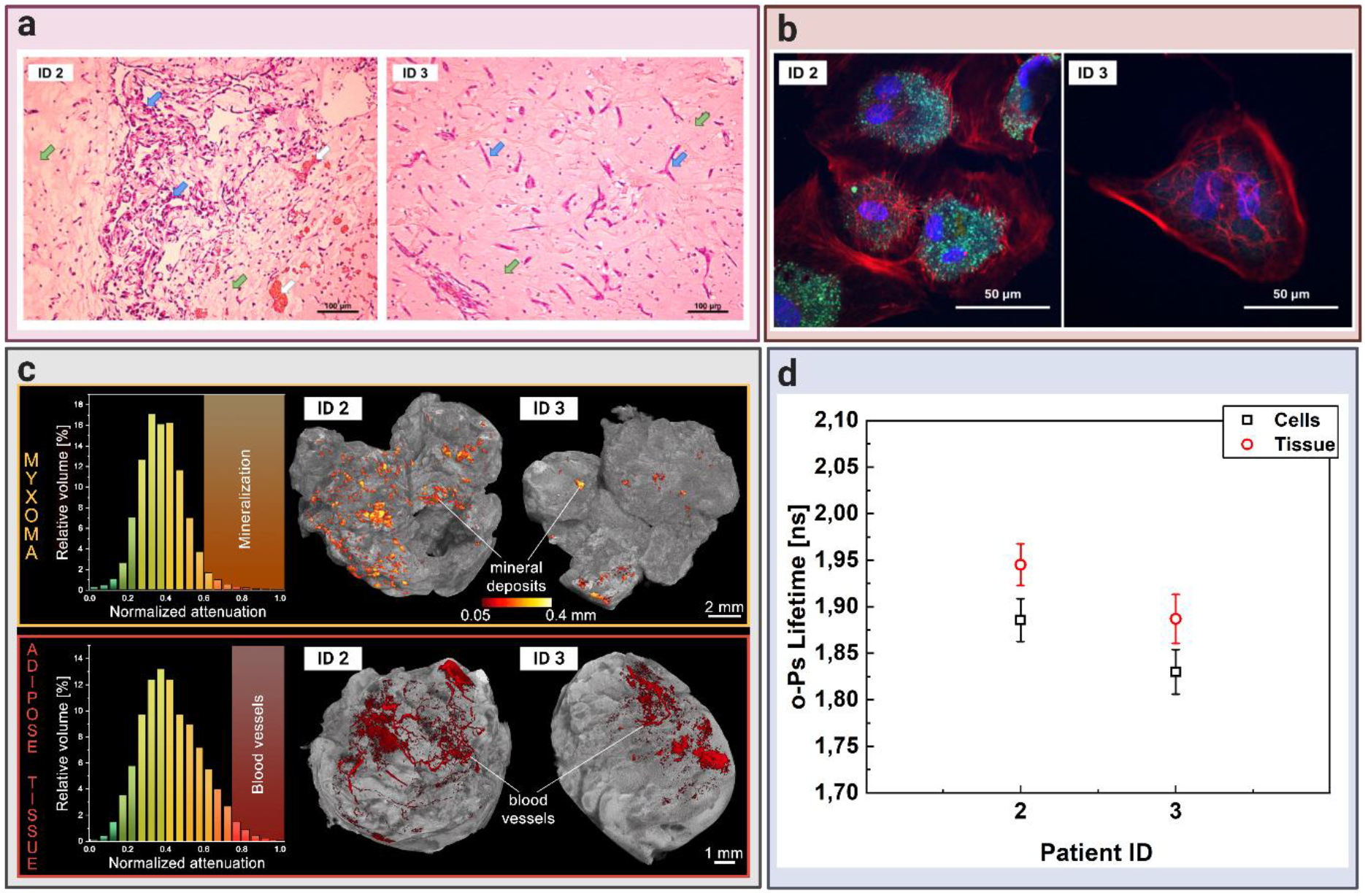
Comparing cardiac myxoma (CM) tissues and the isolated cell line. (A): Micrograph showing exemplary histopathology findings of cardiac myxoma (CM), for patient ID 2 and ID 3 in H&E staining. CM cells stained in purple (blue arrow) can have stellate (ID 3) or globular (ID 2) shape. The red/orange structures (white arrow) correspond to the blood vessels with erythrocytes. The surrounding myxoid matrix is stained in pink (green arrow). (B): Confocal microscopy image with the CM cells stained for F-Actin (red), nucleus (blue), and VE-cadherin (green). The scale bar is 50 μm. (C): Micro-computed tomography results for CM (upper row) and adipose tissues (lower row). Histograms on the left side present normalized X-ray attenuation within the sample: (i) the mineral deposits range from 0.6-1.0 in the CM samples; (ii) the blood vessels range from 0.75-1.0 in the adipose tissue samples. These attenuation ranges have been binarized to extract the mineral deposits and blood vessels for further analysis and visualization. The right side contains volume-rendered 3-D models of the two most representative samples, namely CM and adipose tissues. The internal mineral deposits have been highlighted in the CM model. Its diameter has been color-coded using a heat map. The blood vessels have been colored red in the adipose tissue. (D): Results of the mean ortho-Positronium (o-Ps) lifetime for CM tissue (red circles), isolated from the same patient myxoma cell line (black squares). Patients ID annotations are described in table S1.

Portions of samples used for the positronium investigations were sectioned and irradiated with positrons. We measured the positronium lifetime to compare its value in normal and tumor tissues. In addition, we derived the CM cell culture from two patients to observe the possible difference between the cellular and tissue specimens. Moreover, we conducted X-ray micro-CT imaging on the CM samples for the aforementioned patients to determine the structural heterogeneity of their tumors.

### The mean lifetime of o-Ps atoms in CM is shorter than in adipose tissues

Samples of CM and mediastinal adipose tissue obtained during the surgery were placed in a sterile plastic container, with a cell culture medium supplemented with 10% fetal bovine serum, and antibiotics. We maintained them for the in vitro PALS measurement (Fig. 4A) in the detector system. Each tissue sample was divided into two pieces and interlaced with a radioactive ^22^Na source in Kapton foil. The 22Na radioisotope emits positrons, which may penetrate the tissue up to about 1 mm (*36*), and annihilate with an electron from the biomolecule present in the tissue. The annihilation may proceed directly (e+e-→ photons) or via formation of positronium atom (e+e-→ positronium → photons). Positronium in 25% of cases is produced as short-lived (125 ps) para-positronium (p-Ps) and in 75% cases as long-lived (142 ns) ortho-Positronium (o-Ps). The measurements were performed at room temperature in an aluminum chamber (Fig. 4B) for 1 h to collect 1 x 10^6^ coincidences between annihilation and deexcitation gamma quanta per sample. Fig. 4C presents the exemplary positronium lifetime spectrum obtained for the adipose tissues from the patient ID2. The superimposed lines indicate the distribution of components resulting from p-Ps (green), annihilation in the source material (dark yellow), free positron annihilation (turquoise), o-Ps (blue), and accidental coincidence background (purple). Each component is expressed as

**Fig. 4.**
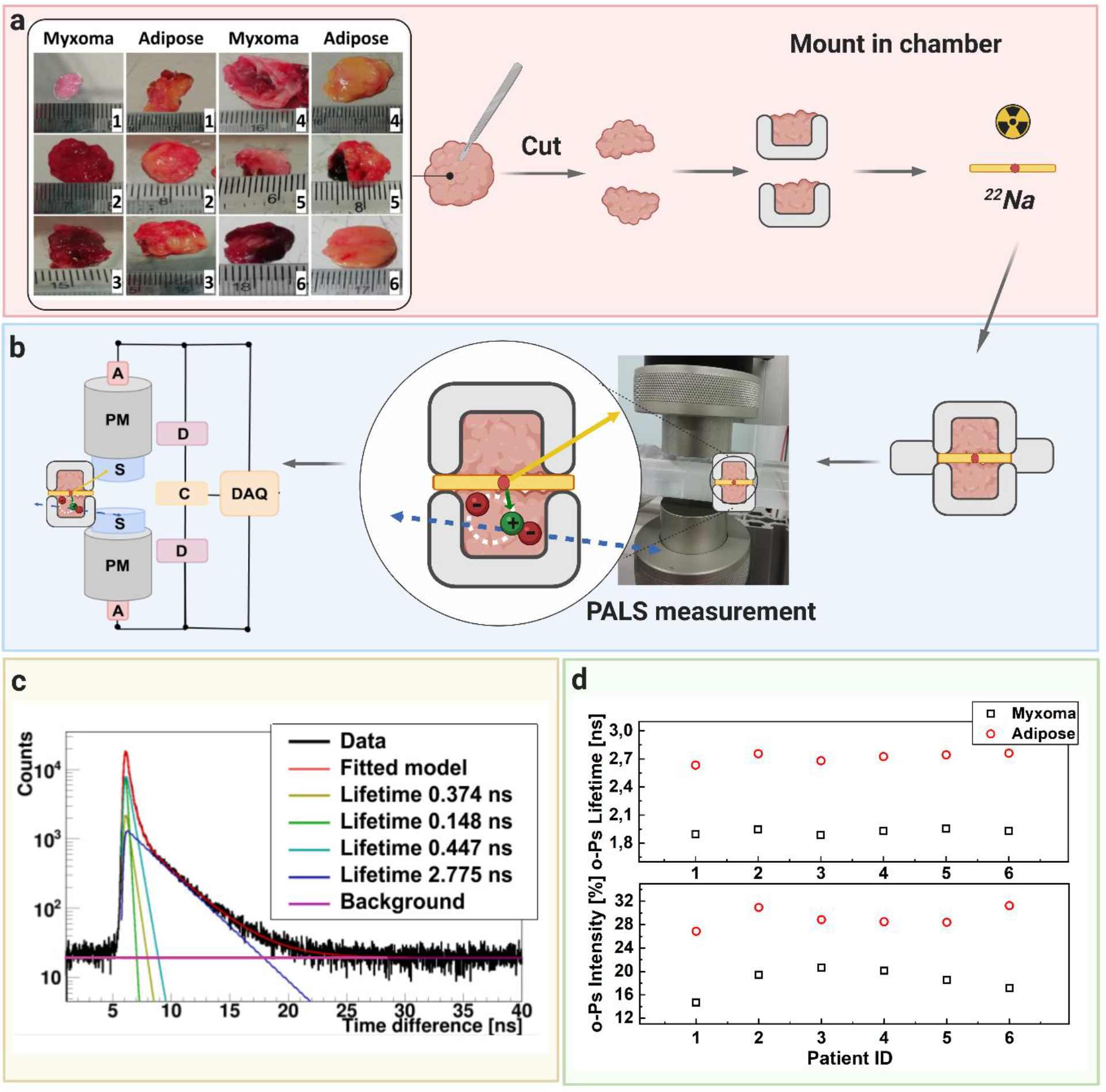
A scheme showing positronium lifetime measurements in cardiac myxoma and adipose tissues. (A): Photographs of non-fixed cardiac myxoma (CM) and adipose tissue samples. The numbers indicate patient ID (Tab.S1). Each sectioned tissue has been cut in halves with a ^**22**^Na radionuclide placed between and inserted into the aluminum measurement chamber. (B): The left part of the panel depicts the scheme of the detection system: scintillators (S), photomultipliers (PM), attenuators (A), discriminators (D), coincidence units (C), digitizer, and a data acquisition system (DAQ). The photograph displays a part of the system together with the plastic rod localized between the scintillators. The superimposed scheme indicates an aluminum chamber inserted inside the rod. ^**22**^Na (red dot) emits (green arrow) positron (+), which annihilates (predominantly into two photons indicated in blue) with electrons (-) in the tissue. Following the positron emission, ^**22**^Na changes into an excited nucleus of ^**22**^Ne, which de-excites almost instantly by the emission of the de-excitation photon (indicated in yellow). The PALS detection system, enables the measurement of the positronium lifetime by registering the time of emission of the de-excitation photon (corresponding to the time of positronium formation) and the time of creating the annihilation photons (corresponding to the time of the positronium decay). (C): For each sample, 1 x 10^6^ coincidences between annihilation and deexcitation *gamma quanta* have been registered, resulting in the lifetime spectrum (example for the adipose tissue - patient ID 2). The analysis enables the extraction of the mean lifetime and intensities of para-Positronium (green line) and ortho-Positronium (blue line) atoms trapped in the intra-molecular voids (*39*). The dark yellow, turquoise, and purple lines denote direct annihilation in the source material, direct annihilation in the sample, and the background because of the accidental coincidences, respectively. The spectra are shifted by the delay coming from detection system configuration (∼5ns). (D): Results of the mean ortho-Positronium (o-Ps) lifetime (upper) and intensity (lower panel) for CM (black squares) and mediastinal adipose (red circle) tissues.

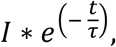

where I and τ denote the intensity and mean lifetime, respectively.

Fig. 4D depicts the mean o-Ps lifetime obtained for the adipose (red circle) and CM (black squares) tissues. For all patients, we observed significant difference between the CM and normal adipose tissue both in o-Ps mean lifetime and intensity. While the averaged mean lifetime and intensity values for CM were 1.92(02) ns and 18.4(2.0)%, respectively, that for the adipose tissue were 2.72(05) ns and 29.1(1.5)%, respectively. The difference in o-Ps mean lifetime and intensity between CM and adipose tissue was 0.8 ns and 10.7%, respectively. In contrast, variations between the patients were around 0.03 ns and 4% in lifetime and intensity, respectively. Thus, there were significant differences in the o-Ps mean lifetime and intensity between normal and tumor tissues. Nonetheless, differences in the o-Ps lifetime and intensity originating from tissue heterogeneity or diversity in the study group did not influence the results.

### Micro-CT imaging and histopathology of CM revealed microscopic differences not influencing the mean o-Ps lifetime

Cardiac CT is commonly used as an additional non-invasive diagnostic modality in cardiovascular diseases, particularly for imaging the aorta and coronary arteries, visualizing the cardiac valves and heart anatomy, and for detecting intravascular and valvular calcifications (*40*). We used micro-CT (enhanced with an iodine contrast agent) as a complementary technique to reveal the structure and composition of CM and adipose tissues. An analysis of the X-ray attenuation distribution within the samples from two representative patients revealed that high attenuation values corresponded with tissue mineralization in CM (normalized attenuation 0.6 - 1.0). A sample obtained from the patient ID2 had greater calcification spots, thus showing that CM tissues were not homogenous in structure. This in turn had been confirmed by the histopathology examination. The aforementioned patient had globular shaped CM cells and vascularization spots (Fig. 3A). In cell culture, the CM cells isolated from the patient ID2 had a typical mesenchymal morphology and expressed the endothelial binding protein: vascular endothelial (VE)-cadherin (*31*). In contrast, the CM sample from the patient ID3 had relatively fewer calcification spots. Moreover, the cells appeared to be stellate-like in histopathology. CM cells from this patient did not show VE-expression in cell culture (Fig. 4B). In addition, vascularization in the adipose tissues was represented and visualized for both patients showing different attenuation values than for calcification (normalized attenuation 0.75-1.0) (Fig. 3C). We did not detect blood vessels in CM. Similarly, there were no mineral deposits in the adipose tissues. There is a significant diversity in the spatial distribution of highly attenuating regions in the samples, particularly in the dispersion of mineral deposits.

In contrast to the calcification deposits revealed by micro-CT, microvessels cannot be visualized in CM with the above-mentioned technique. This necessitates a higher resolution approach. The use of positronium imaging as a complementary method to CT or MRI can distinguish the histological changes and structural differences on a nanometer level.

### Comparing the CM tissue and the isolated cell line reveals similar o-Ps lifetime characteristics

We established the myxoma cell culture from tissues operated from two patients, namely ID2 and ID3. This helped us compare the mean o-Ps lifetime in the CM cells and CM tissues, which comprise a milieu of extracellular matrix and blood vessels, in addition to cells. The Methods Section in Figure 5a outlines the workflow of CM cell isolation, culture, and sample preparation for the positronium lifetime measurement. Furthermore, it also presents micrographs of cell cultures upon seeding (0h), 24 h later, and the secondary culture. We observed the floating CM cells and the number of erythrocytes while seeding. The CM cells attached to the surface of the culture dish surface 24 h later. The remaining debris and erythrocytes were washed out. After a week following the isolation, the CM cells developed a mesenchymal-like shape having multiple nuclei, mostly around 2-3 per cell in the secondary culture. The myxoma cells had a diameter of roughly 40-50 µm (Fig. 3B).

For the positronium lifetime analysis, the CM cells were harvested from the cell culture, sedimented to obtain roughly 20 million cells (Figure 5a), and placed in an aluminum chamber at 37°C for 1 h (Fig. 3B, fig. S1B). We determined the cell viability before and after the measurement and obtained a difference of 10.0(2.3)%. The exemplary positron lifetime spectra for the measured samples are presented in the Methods Section in Fig. S1B.

The determined mean o-Ps lifetime for the cultured CM cells (red circles) from both patients (ID2 and ID3) was lower than those for the raw CM tissues (black squares): 1.89(3) ns vs 1.95(1) ns and 1.83(3) ns vs. 1.89(3) ns, respectively (Fig. 4D). These results are present within 2 standard deviations. Despite removing the surrounding tissue matrix from the myxoma sample, such as extracellular proteins and vessels, the tendency to obtain a higher signal for the patient ID2 was preserved. CM cells from the patient ID2 expressed greater adhesion proteins (for example, VE-cadherins), which may be related with different molecular structures and metabolism, and influence the o-Ps lifetime (Fig. 4B).

## Discussion

Methods used to confirm a CM diagnosis varied in past decades. Earlier, patients were diagnosed by cardiac catheterization. However, TTE is performed routinely in recent times, with MRI or CT scan imaging being used as a complementary diagnostic modality (*25-27,40,41*). TTE can be extended by a transesophageal approach that facilitates an intraoperative cardiac investigation to confirm tumor localization (*27,40*).

In practice, echocardiography can usually distinguish between the three intracardiac masses, namely tumor, thrombus, and vegetation, with an accuracy 80% (*41-43)*. A presumptive TTE diagnosis can be made for primary cardiac neoplasm, including CM, for some distinctive features (*21*). Therefore, MRI and CT are constantly needed to improve the definitive diagnosis. Nonetheless, these techniques provide limited tissue characterization.

The recent advent of high sensitivity total-body PET (*1,3,4,7*) and the invention of positronium imaging offer new perspectives for improved diagnostic and prognostic assessment, based on cell and its *milieu* alterations at the molecular level (*2,11–14,44*). Positronium is copiously produced in the intra-molecular spaces during routine PET imaging. It can be effectively used as a diagnostic parameter, complementary to the standardized uptake value image (*2,37*). We demonstrated for the first time substantial differences in the positronium mean lifetime and the production probability for cardiac neoplasm (myxoma), compared to normal mediastinal adipose tissues. Thus, our findings offer promising prospects for improving the diagnosis of heart diseases using positronium imaging. Notably, the determined differences of the mean positronium lifetime in CM and adipose tissues are at the level of more than 10 standard deviations for each examined patient. However, the mean o-Ps lifetime values in CM were constant within 2 standard deviations in the study group of patients.

PET is currently used as a standard in cardiology for myocardial perfusion evaluation (*16,32–35*). Therefore, the new functionality of PET devices that enables positronium imaging, simultaneously with standard SUV imaging, shall constitute a natural enhancement of the diagnosis by preserving the current PET imaging protocol.

We used sodium ^22^Na isotope as an emitter of positrons (e+), considering the role of sodium fluoride in cardiovascular imaging (*32,35*). To date, researchers have applied ^18^F-sodium fluoride only for PET imaging, with ^18^F radionuclide as a positron emitter. However, labeling sodium fluoride with ^22^Na (^22^NaF) enabled simultaneous SUV and positronium imaging (*1*). This can be attributed to the prompt photon emission in ^22^Na decay. Another example of a radiopharmaceutical applicable for the positronium imaging of cardiovascular system is the Food and Drug Administration–approved ^82^Rb-Chloride. This in turn would facilitate assessing the perfusion of the cardiovascular system (*1,45*). Similar to ^22^Na, ^82^Rb isotope also emits a prompt photon, in addition to a positron. Sitarz et al. have reported on prompt photon positron emitters that look promising for PET imaging (*46*). Concurrent application of the β+γ emitter with a novel radiotracer, e.g. the ligand of serine protease fibroblast activation protein (FAP), which is a biomarker of activated (myo)fibroblasts, will give a new advantage in assessment of post-myocardial infarction remodeling (*47*). Moreover, examples of tracers proposed for PET imaging with the aforementioned isotopes are summarized in the reference by Moskal and Stępień (*1*).

In summary our results demonstrated the usefulness of positronium (its lifetime and intensity) to differentiate between pathologies, that is, benign cardiac tumors and adipose tissues. These data evaluate an o-Ps lifetime as a new biomarker for cardiovascular imaging, particularly for imaging intracardiac masses to reveal their structural composition. Structural heterogeneity is characteristic of CM tumors. However, we showed that the macroscopic (morphology) and microscopic (micro-CT) structural changes did not influence the value of the observed parameter (mean o-Ps lifetime), which reflected changes on a molecular (nanoscale) level.

## Acknowledgments

We acknowledge the support and help of Dr. L. Rudnicka-Sosin MD for the histopathological examination. We are thankful to Prof. B. Jasińska and Dr. M. Gorgol for their support in the preparation of radioactive sources, and to W. Migdał and A. Wróbel for their technical support. We acknowledge the help and support of Prof. Z. Rajfur and T. Kołodziej for enabling and conducting the confocal microscopy imaging. We acknowledge the financial support by the Foundation for Polish Science through the TEAM POIR.04.04.00-00-4204/17 Programme, the National Science Centre through grants No. 2017/25/N/NZ1/00861 and 2019/35/B/ST2/03562, the Ministry for Science and Higher Education through grants No. 7150/E-338/SPUB/2017/1, K/ZDS/007227, N41/DBS/000230, and CRP/0641.221.2020, and the SciMat Priority Research Area budget under the program Excellence Initiative Research University at the Jagiellonian University.

## Funding

Foundation For Polish Science TEAM POIR.04.04.00-00-4204/17 (PM)

National Science Centre of Poland 2019/35/B/ST2/03562 (PM)

National Science Centre of Poland 2017/25/N/NZ1/00861 (EK)

Ministry for Science and Higher Education 7150/E-338/SPUB/2017/1 (PM)

Ministry for Science and Higher Education K/ZDS/007227 (GG)

Ministry for Science and Higher Education N41/DBS/000230 (GG)

Ministry for Science and Higher Education CRP/0641.221.2020 (PM, ES)

SciMat Priority Research Area Excellence Initiative - Research University at Jagiellonian University (EK, KD)

## Author contributions

Conception and design of the work: E.Ł.S, P.M

Data collection: B.L., E.C., E.K., K.D., S. N.

Software: K.D.

Hardware: E.C., S.N

Patients’ enrolment and data curation: G.G. Visualization: B.L. and E.K.

Data analysis and interpretation: B.L., E.C., E.Ł.S., E.K, G.G., K.D., P.M., S. N.

Funding acquisition: E.K, E.Ł.S., G.G., P.M.

Project administration: E.Ł.S, P.M

Drafting the article: E.Ł.S., E.K., P.M.

Critical revision of the article: E.Ł.S., P.M.

Final approval of the version to be published: B.L., E.C., E.Ł.S., E.K, G.G., K.D., P.M., S. N.

## Data and materials availability

All data are available in the main text or the supplementary materials.

## Supplementary Materials

### Materials and Methods

#### Study group characteristics

We analyzed and compared non-fixed myxoma samples with adipose tissues obtained from the patients. Each patient completed a screening interview, laboratory test (blood counts and metabolic panel), electrocardiogram, and urine toxicology. In all cases, the patients were diagnosed by TTE. Therefore, a histopathological examination of the specimen postoperatively confirmed the structure type. Table S1 summarizes information on the patient’s health, type of tumor, and other demographic factors. This study was approved by the Bioethical Commission of the Jagiellonian University (approval number 1072.6120.123.2017) and an informed consent was obtained from each patient.

#### Tissue sample preparation

During the surgery, the extracted tumor was aseptically sectioned into two pieces. While one piece was sent for the histopathology examination, the other one was placed in a sterile plastic container filled with Dulbecco’s Modified Eagle’s medium (DMEM) cell culture medium, high glucose (Cat. No. 61965026, Gibco®, Paisley, UK) with 10% fetal bovine serum (Heat Inactivated, Brazil Origin, Cat. No. 10500064, Gibco®, Paisley, UK), penicillin (10.000 U/mL), streptomycin (10.000 µg/mL) (Cat. No. 15140122 Gibco®, Paisley, UK), and amphotericin B (25 µg/mL) (Cat. No. 15290026, Gibco®, Paisley, UK). It was transported to the laboratory for the positron annihilation lifetime spectroscopy (PALS) examination within 4 h of extraction. Both myxoma and adipose tissues were sectioned into two pieces. Each PALS-measured sample consisted of two parts with the ^22^Na source in between (Fig. 3B). After PALS measurement tissues were fixed in 4% formaldehyde in phosphate buffered saline (PBS) for micro-CT imagining.

#### Histopathology examination

We conducted histopathological examinations for all extracted tumors. Robust samples were fixed in 4% formaldehyde in phosphate buffered saline (PBS), embedded in paraffin. The routine histopathology procedure was performed with H&E staining. The examination revealed elongated, round, or stellate single cells and tenuous cords, typically infiltrated by lymphocytes and macrophages, and surrounded by myxoid stroma.

#### Cell Culture

We derived myxoma cell cultures from two patients. The tissues were aseptically dissected with scalpel. We incubated the portion of tissue designated for cell isolation in a petri dish for 48 h. It comprised DMEM medium, high glucose (Cat. No. 61965026 Gibco™ Paisley, UK) with 10% fetal bovine serum (Cat. No. 10500064 Gibco™ Paisley, UK), penicillin (100 U/mL), streptomycin (100 µg/mL) (Cat. No. 15140122 Gibco™), amphotericin B (0.25 µg/mL) (Cat. No. 15290026 Gibco™), L-Glutamine (2 M) (Cat. No. 25030081 Gibco™ Paisley, UK), and collagenase II (200 U/mL) (Cat. No. 17101015 Gibco™, Paisley, UK). Following incubation, the tissue with the medium was squeezed through a 70 nm nylon mesh to isolate cells from the extracellular matrix. It was centrifuged at 260 g for 10 min, seeded on a T75 cm^2^ cell culture dish, and cultured at 37°C and 5% CO2 atmosphere. Myxoma cells started attaching to the bottom of the dish after 24 h. We regularly washed the cells with PBS w/o Ca^2+,^ Mg^2+^ (Cat. No. 10010015 Gibco™ Paisley, UK) during the first week of culture. This helped us wash out the erythrocytes. The cells were then incubated with 0.25% Trypsin-ethylenediamine tetra-acetic acid (Cat. No. 25200072 Gibco™ Paisley, UK) for 10 min, and removed from the flask with a cell scraper. We then centrifuged the cells in 260 g for 10 min. Following a spin, the cells were counted with Trypan Blue dye by an Automatic Cell Counter LUNA II, and seeded to the flasks at an appropriate density. Myxoma cells grow slowly and are usually seeded at a density of 2.5 x105 cell/cm^2^. They reach 80% confluence after 50 days. We prepared the cell samples for PALS measurement by passaging in a standard way, followed by counting the cells and suspending the pellet in a medium. It was later centrifuged in 500 g for 90 s. The supernatant was discarded thoroughly, and the cell pellet was transferred to a measurement chamber with a spatula. We determined the viability before and after each measurement to confirm that the setup conditions and cell handling were appropriate for the culture, and the high percentage of dead cells did not influence the results.

#### Microscopy imaging

We regularly checked the cell morphology by means of a Nikon Eclipse TS100 inverted optical microscope. The photographs were captured with a Nikon DS-Fi1c camera and are presented in Figure 3a. We conducted confocal microscopy studies for the cardiac myxoma cell culture on Zeiss Axio Observer Z.1 with an LSM 710 confocal module with Alexa FluorTM 647 Phalloidin (Cat. No. S32357 Invitrogen TM Paisley, UK) and the DAPI-stained (Cat. No. 62248 PierceTM Paisley, UK) for F-Actin and nucleus, respectively. VE-cadherin (BV9) Antibody (Cat. No. sc-52751, Santa Cruz Biotechnology, Inc.’s, US) and goat anti-mouse IgG-FIT secondary antibody (Cat. No. sc-2010, Santa Cruz Biotechnology, Inc.’s, US) with fluorescein isothiocyanate were used for the immunofluorescence staining. The images were processed with ZEN lite software.

#### Positron Annihilation Lifetime Spectroscopy

The positron annihilation lifetime spectrometer presented in Figure 3b constituted two H3378-51 Hammamatsu photomultipliers, equipped with BaF2 cylindrical scintillators, with a diameter and height of 38 mm and 25 mm, respectively. It had been manufactured by Scionix. Detectors were powered by CAEN SY4527 high voltage power supply. Signals from the photomultipliers were attenuated by 10 dB (indicated as A) and delivered to LeCroy 608C constant fraction discriminator (D). Different thresholds were applied to the signals from diverse detectors in the latter. Considering the varying thresholds set on the discriminator, while one detector was considered as ‘START’ (registers 1274 keV gamma quanta), the other one was considered as ‘STOP’ (registers 511 keV gamma quanta). We set the coincidence time window to 110 ns on the LeCroy 622 coincidence module. The data was acquired by a digitized DRS4 evaluation board. We used radioactive 22Na source with an activity of 1 MBq, sealed between 6 µm thick Kapton foils for all measurements. The samples were measured in an aluminum chamber with a source sandwiched between them. They were placed in a temperature-controlled holder. While the tissue samples were measured at room temperature, the cell culture samples were measured at 37 °C.

#### Micro-CT

We used Lugol staining solution to enhance the X-ray attenuation within the samples (*48*). The samples were fixed with 4% formaldehyde and placed in an individual dish containing 50 mL Lugol solution (I3K) (Cat. No. 62650 Sigma-Aldrich). They were stored for 5 days (4° C). Before scanning, each sample was washed with saline solution and placed in individually designed 3-D printed sample holder to prevent motion during the scanning. We conducted the micro-CT investigation with Bruker SkyScan 1172 (Kontich, Belgium) scanner. The X-ray energy was set to 80 keV. No physical filter had been used. We captured the images with a pixel size of 8.97 µm. Each projection image was averaged out of eight frames to enhance the signal-to-noise ratio. Image reconstruction was performed using NRecon version 1.7.3.1 software by Bruker Micro-CT (Kontich, Belgium). The cross-section images were processed and analyzed using CTAnalyser 1.20.3.0 software by Bruker Micro-CT (Kontich, Belgium). The 3-D volume rendered models were prepared using CTVox 3.3.0 software by Bruker Micro-CT (Kontich, Belgium).

#### Statistical Analyses

Results of PALS are presented as the mean o-Ps lifetime and intensity. They were analyzed by the PALS Avalanche program, developed by the Jagiellonian-PET collaboration (*39,49*). Uncertainties of each measured point are a statistical error of the fitted function. The viability test and cell count were analyzed by means of the LUNA-II™ Automated Cell Counter (Logos Biosystems).

---

**Figure S1.**
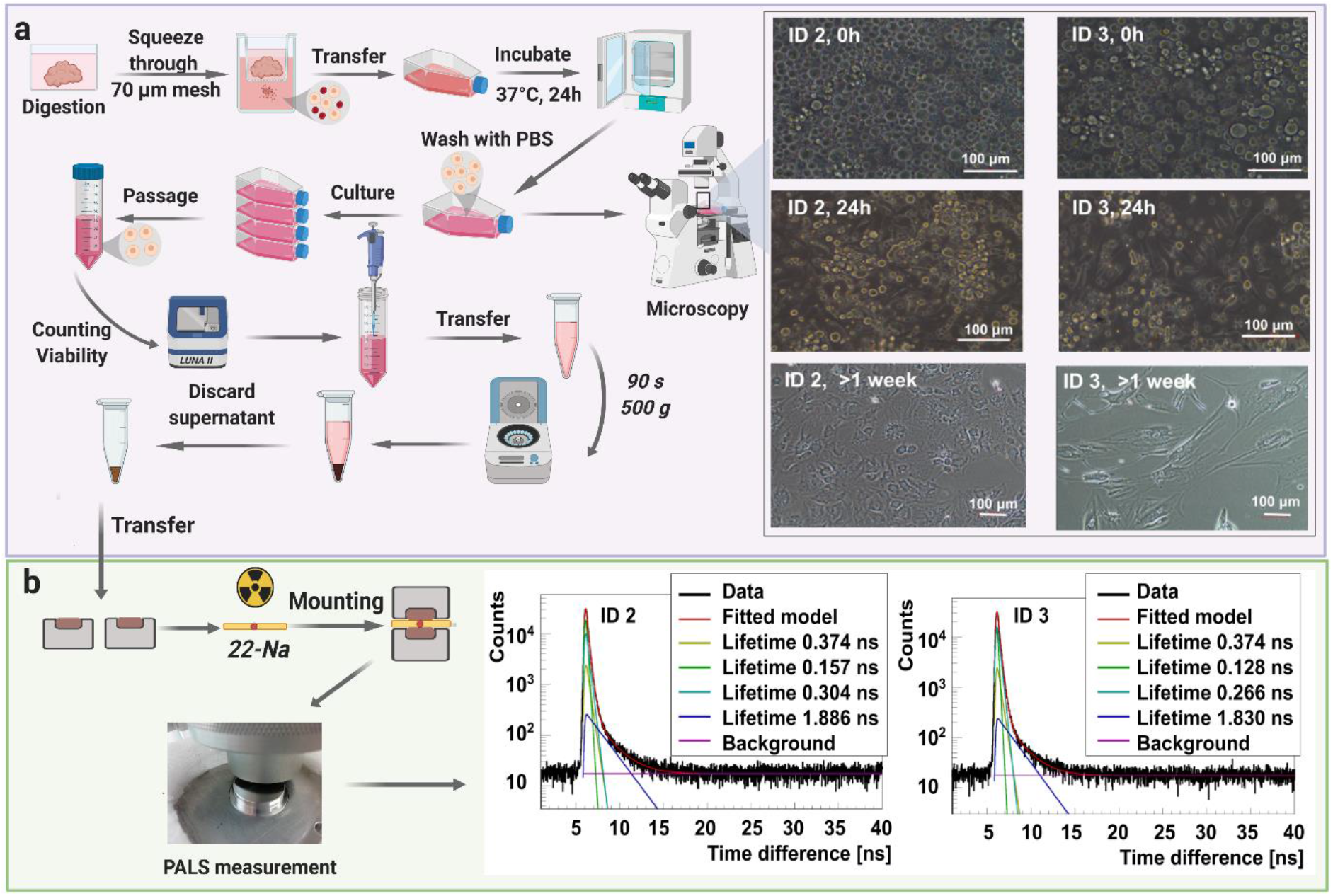
Myxoma cell culture isolated from the tumor of patient ID 2 and ID 3. (A): (left) The workflow of myxoma cell culture isolation. Cells have been isolated by tissue digestion and cultured to obtain the highest possible number of cells for the experiment. A micrograph presenting the primary culture of cells isolated from cardiac myxoma, upon seeding, and 24 h later with erythrocyte contamination. The latter has been washed out during the primary culture. Secondary culture after the 1st passage has been established after 1 week. A scale bar of 100 μm. (B): (left) The workflow of PALS measurement. The centrifuged cells have been placed in both parts of the aluminum chamber with a radioactive source (red dot) encapsulated between them. The chamber has been mounted between detectors in the temperature-controlled aluminum holder. The measurements have been performed at 37 °C. (right) Positronium lifetime spectra with fitted components for the myxoma cell culture isolated from the given patient. The dark yellow, green, turquoise, and blue lines denote direct annihilation in the source material, p-Ps annihilation component, direct annihilation in the sample, and o-Ps annihilation component, respectively. The spectra are shifted by the offset coming from the detection system configuration (∼5ns). The patient ID annotations are described in table S1.

**Table S1.** Clinical characteristic of a study group.

| Patient ID | Sex | Age | Tumor localization | Tumor type | Tumor size (cm) | Metabolic disease | Cardiovascular disease | Diabetes |
| --- | --- | --- | --- | --- | --- | --- | --- | --- |
| 1 | F | 78 | Left atrium | solid | 3x3 | hyperlipidemia | hypertension, Ischemic heart disease, patent foramen ovale | - |
| 2 | M | 57 | Left atrium | papillary | 8x3 | extreme obesity | paroxysmal atrial fibrillation, hypertension | - |
| 3 | F | 59 | Left atrium | papillary | 6x5 | - | hypertension, ischemic heart disease | Type II |
| 4 | F | 84 | Left atrium | solid | 3.5x2.5 | gout, hypercholesterolemia | hypertension, paroxysmal atrial fibrillation, varicose veins, ischemic heart disease | - |
| 5 | F | 52 | Left atrium | solid | 3x2 | - | - | - |
| 6 | M | 64 | Left atrium | solid | 2x2 | hypercholesterolemia<br>chronic renal failure | fixed atrial fibrillation, peripheral atherosclerosis, congestive heart failure | - |

